# Unveiling the Emotional Turmoil: How Covid-19 impacted researchers and the pursuit of emotional well-being in academia

**DOI:** 10.1101/2023.11.15.567306

**Authors:** Crista Weise, Nuria Suñe-Soler, Mariona Corcelles, Anna Sala-Bubaré, Monsterrat Castelló

## Abstract

The Covid-19 crisis unprecedentedly required researchers to adapt to significant changes in their work and personal lives. Our study aims to fill this gap analysing the Covid-19 emotional impact and confinement potential disruptions on researchers’ activity (specifically, those related to working conditions, caring responsibilities, health, balance, and social support) considering the modulating role played by age, gender, and job position. An online survey was distributed during the first lockdown period of the Covid-19 pandemic, and answers from 1301 researchers (ECR %, senior researchers %) working in Sciences (28.1%), Social Sciences (25.9%), Humanities (16.2%), Health (16.2%) and in Engineering and Architecture (13.5%) were collected. The study highlights that the initial lockdown during the Covid-19 pandemic had a significant emotional impact on researchers, exacerbating pre-existing emotional distress and burnout within this group. Factors such as age, health, gender, and difficulties in balancing work and family life were associated with an increased risk of burnout and emotional distress. Lack of social support was identified as a significant risk factor, while the academic culture prioritizing productivity over well-being contributed to the issue. These findings underscore the need for greater support and cultural changes in academia to preserve researchers’ mental health and prevent the chronicization of mental health issues in young academics.

## Introduction

The Covid-19 crisis unprecedentedly required researchers to adapt to significant changes in their work and personal lives, at a time when science was called to respond quickly with effective solutions to address the major effects of the pandemic. This tension arose in the first quarter of 2020, in a situation of confinement that many of us had never experienced before, with great uncertainty about the future, limited agency, strict regulations of the way we socially interact, potentially unsuitable work environments, and/or increased responsibilities and concerns related to health and personal life.

In regular conditions, academic career development involves significant challenges and is strongly impacted by research conditions, institutional and personal factors (Skakni & McAlpine, 2017). It requires coping with emotionally-taxing situations within a scientific and disciplinary community (Cotterall, 2013) usually affecting work performance and personal life (Kurtz-Costes et al., 2006; Stubb et al., 2011).

Staff working in academia has been considered high risk population in terms of mental health and well-being at work. Levels of burnout appear higher among academics than in general work populations. Large proportions of graduate students’ report having symptoms of depression, emotional problems, or problems related to high levels of stress (Guthrie et al., 2018).

The Covid-19 pandemic added to this situation several disruptions (Kurtz-Costes et al., 2006) modifying the work and life environment, encompassing new and different challenges, and increasing the stress over academic tasks. Among those disruptions, research has highlighted the impossibility of accessing the logistical structures necessary to carry out the research activity, the slowing down or temporary suspension of research events, long delays in producing scientific outputs, and/or difficulties in collecting data and disseminating research findings (Aczel et al., 2021; Gilmartin et al., 2021; Rodrigues et al., 2021; Sarah et al., 2021). Researchers had to deal with those disruptions with decreased levels of institutional and social support (Rupnow et al., 2020). Research centres or universities had to decide which essential departments and administrative services would continue to operate virtually, which significantly shortened the various supports provided to researchers

Confinement also reduced informal contact with colleagues and increased researchers’ sense of social isolation and perception of loneliness (Aczel et al., 2021; Gilmartin et al., 2021; Sandhu et al., 2021; Sarah et al., 2021; Wang & DeLaquil, 2020), which, in turn, was associated with higher levels of stress, anxiety, and burnout (Mäkiniemi et al., 2021; Rotărescu et al., 2020; Suart, Neuman & Truant, 2022). Furthermore, during the confinement many of the grant competitions this population rely on were postponed or cancelled (Kent et al., 2020; Rodrigues et al., 2021). Some researchers reported not receiving new funding to restart research projects and an increased level of uncertainty about career future, with some experiencing the prospect of even losing their jobs (Byrom, 2020; Sandhu et al., 2021; Sarah et al., 2021; Tay, 2020). Additionally, training, networking, and job opportunities had also decreased during the confinement, leaving those in the early stages of their career with the perception of fewer options (Creaton & Handforth, 2021; Jamali et al., 2023).

Another aspect research has signalled as particularly challenging during confinement was assuring work-life balance, an area already problematic for academics and researchers prior to the pandemic (Badri, 2019). Previous studies found that a positive perception of work-life balance favours better performance as teachers, researchers and administrators, and that, on the contrary, overload and stress caused by work-family tension entail risks for individual health, increase depression and decrease productivity (Duxbury et al., 1994). The switch to an online mode of working during the Covid-19, have blurred the traditional boundaries between work and personal life. In addition to the loss of structure, working at home reduced social contact with colleagues, leading to increased stress and, for many researchers, difficulties in balancing research and family responsibilities.

Additional time spent caring for children and/or family members impaired opportunities to work on research (ISSCR, 2020). Confinement required additional effort to reconcile different professional roles and tasks, especially for teaching researchers who had to hastily move learning activities to a virtual environment with additional workload for learning new computer skills, adapting strategies and materials, and orienting students (Biswakarma et al., 2021; Johnson et al., 2020; Rodrigues et al., 2021; Sahu, 2020).

The consequences of these disruptions disproportionately affected certain groups of researchers. The highest negative impacts on research dedication or productivity were found among researchers performing laboratory-based or field work (Creaton & Handforth, 2021; Myers et al., 2020; Omary et al., 2020), those with precarious positions (Byrom, 2020; Tay, 2020; Sandhu et al., 2021; Sarah et al., 2021; Suart, Neuman & Truant, 2022), female researchers, and those with children or other caring responsibilities (Cardel et al., 2020; Gabster et al., 2020; Garrido et al., 2021; Myers et al., 2020; Pereira, 2021). Although some studies did not find parental status to be related to productivity (Cole & Zuckerman, 1987; Sonnert & Holton, 1996), other suggest that the experience of balancing family and career may be somewhat different for women than for men, especially the pre-tenured, who have worst work conditions (Milkie & Peltola, 1999; Profeta, 2020).

On the other hand, within the crises are the seeds of opportunities. In all the studies reviewed, we detected a considerable percentage of researchers who reported insignificant negative effects of confinement on their activity and some of them even recognized that it contained opportunities or positive benefits. For this group of researchers, confinement offered conditions to develop scientific activities involving concentration (e.g., reading the literature, writing and analysing data), more time to study and plan future research activities. They also found the potential benefits of increased digital collaboration, reduced energy footprint and environmental impact to be relevant (Center N.D., 2020; Sarah et al., 2021). Likewise, some researchers reported that the online work greater flexibility helped them improve work-life balance (Aczel et al., 2021).

The differences in personal appreciations of the constraints or opportunities that confinement entailed could be associated with some variability regarding its emotional impact. At a general level, the confinement has been related to psychological strain and exhaustion with the subsequent loss of focus and increased anxiety on this population (Byrom, 2020; Centre, 2020; ISSCR, 2020; Kent et al., 2020; Rodrigues et al., 2021; Sachini et al., 2021; Sahu, 2020; Suart et al., 2021; Suart, Neuman & Truant, 2022). Again, these studies results are heterogeneous, with a non-negligible number of researchers reporting no significant emotional impact. This variability could be explained by the differences among the idiosyncratic challenges researchers faced, associated not only with the potential disruptions caused by the confinement but also with systematic barriers that traditionally have been affecting certain groups of researchers (e.g., female or early-career researchers).

Some studies have pointed to the influence of family climate and family functioning as factors that clearly influence people’s mental health; mindfulness and psychologically flexible response styles (as opposed to rigid and avoidant response styles), are behavioural repertoires that have previously been shown to buffer the impact of stress and facilitate well-being (Gloster et al., 2020).

However, except for a few studies focused on ECRs (Byrom, 2020; Mäkiniemi et al., 2021; Suart et al., 2021), little is still known about the main predictors of higher levels of emotional distress on this population during confinement. Our study aims to fill this gap analysing the Covid-19 emotional impact and confinement potential disruptions on researchers’ activity (specifically, those related to working conditions, caring responsibilities, health, balance, and social support) considering the modulating role played by age, gender, and job position. Discriminating the particularities of the confinement emotional cost according to the researchers’ idiosyncratic experiences is critical to offer tailored support to this community and effectively protect their well-being in potentially future similar situations. Considering that before the pandemic there were already concerns about the levels of psychosocial well-being of this population (Bekkouche et al., 2021; Guthrie et al., 2018) adds even more relevance to this analysis.

Two specific objectives were established to fulfil the study general aim:

(1) To analyse the Covid 19 first confinement emotional impact on researchers’ activity
(2) To determine the modulating effects on the emotional distress of variables such as age, gender, health, research position, working conditions, caring responsibilities, health, balance, and social support

## Method

The final sample includes responses from 1,301 Catalan^i^ researchers. The participants (mean age = 43.85; SD= 12.436) were evenly distributed within the study’s essential variables. Regarding gender, 53.9% were women, 45.1% men, and 1% identified with non-binary gender categories. In terms of discipline, 28.1% were doing research in Sciences, 25.9% in Social Sciences, 16.3% in Humanities, 16.2% in Health Sciences, and 13.5% in Engineering and Architecture. Concerning the career stage, 29.6% were doctoral students, and 70.4% were doctor researchers, relatively well distributed throughout the different stages (12.3% postdocs, up to 5 years of experience; 21.4% mid-career, between 6-15 years of experience; and 34.7% senior researchers, more than 15 years of experience). Regarding their personal situation, 17.06% of researchers stated that they had suffered from a health problem during the lockdown and 45.64% took care of other people (e. g., children, people with some dependency).

We designed a questionnaire^ii^ to collect data on the impact of Covid-19 lockdown on research activity, career, and wellbeing of active researchers in Catalan-speaking communities. The data were collected from The questionnaire contained 25 items organised in two first sections devoted to collect sociodemographic data and working conditions and other five sections looking for the Covid-19 lockdown effects on emotional wellbeing, research activity, social perception of science and social support. In this study, we used the variables gender, age, and research position and the dimensions of emotional wellbeing, social support, working conditions, balance, caregiving, and health. All questions were closed-ended, with the option to specify other answers that were subsequently re-coded (see Table 1).

**Table 1.**
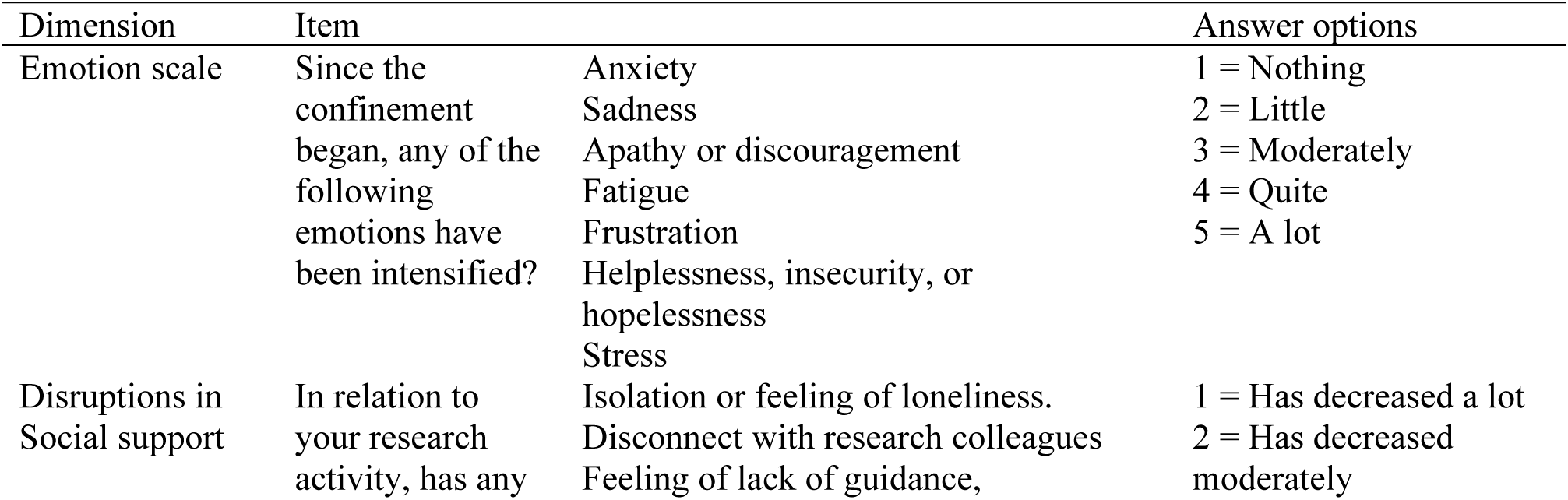
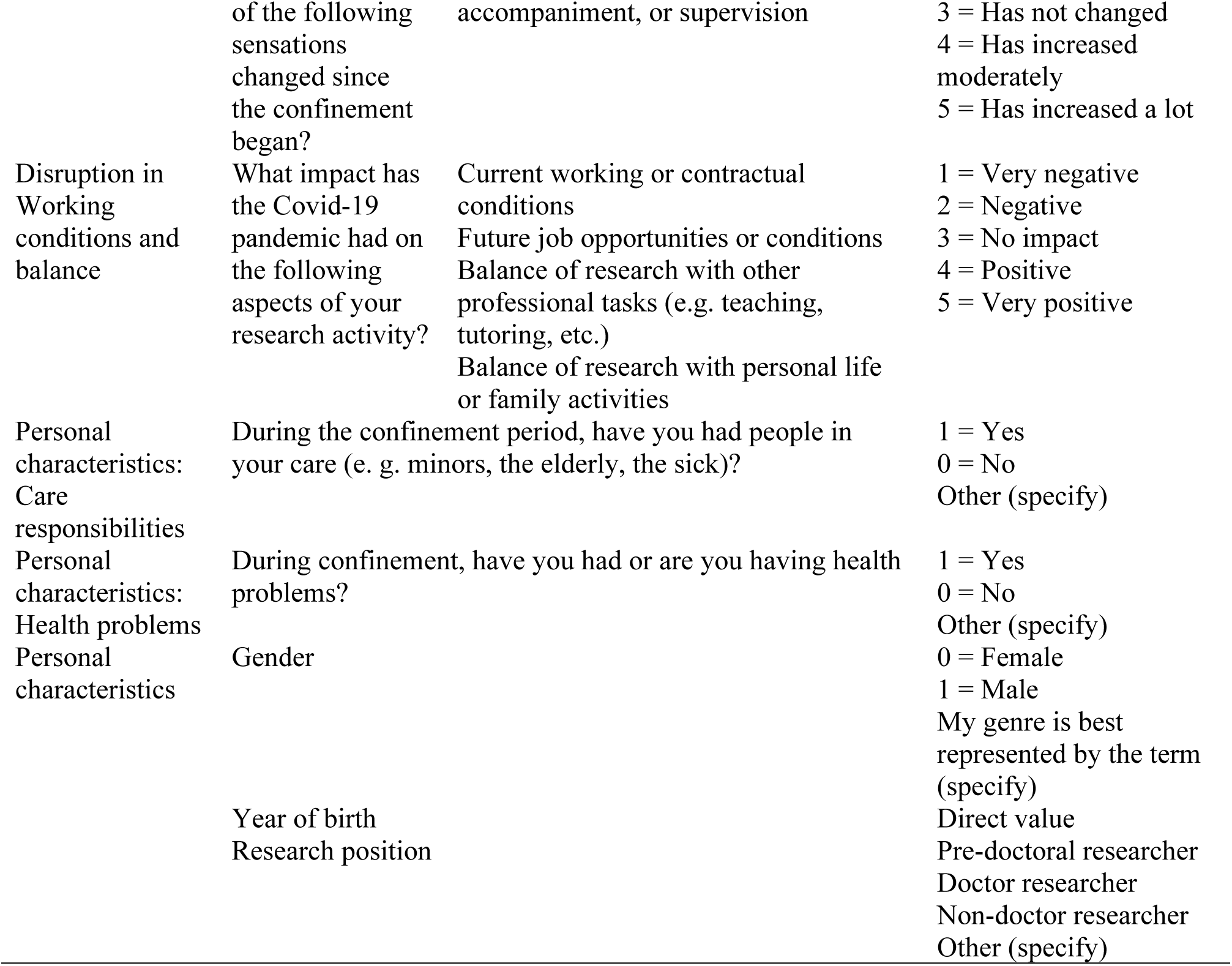
Dimensions and items of the questionnaire “Covid-19 pandemic and research activity” used in this study.

The procedure for contacting participants was based on an automated recruitment method of public emails, approved by the ethics committee of our university and the data protection authority of our country. An algorithm was designed to build a distribution list with the professional email addresses of research staff from all the main universities in the Catalan-speaking regions (i.e., Catalonia, Balearic Islands, and Valencia). From May 8 to June 30, 2020, a single email was sent to potential participants with explanatory information about the project. The link to the questionnaire was not included in the email to increase transparency and ensure participants had complete information about the project characteristics and the ethical treatment of data before answering.

Before proceeding to answer the questionnaire, through an additional question on the form, the participants explicitly accepted by written the informed consent, guaranteeing the trustworthiness of the participants. All participants enrolled voluntarily, with the possibility of leaving the study at any time they considered appropriate.

The data were deposited in a secure server approved by the university. All data were anonymized guaranteeing the confidentiality and ethical treatment of the data. The Declaration of Helsinki was properly followed as express in the ethics committee document that guarantees its compliance (Ref. APR-FPCEE2021/02).

To analyse data, we first conducted descriptive statistics on all the variables. Then, maximum-likelihood Exploratory Factor Analysis with oblimin rotation was carried out to identify the emotional scale underlying factors. We verified data was suited for factor analysis, as Kaiser-Meyer-Olkin values were higher than 0.83 and Bartlett’s test was significant (X2 (21) = 4251.411, p < .001). All assumptions (Yong & Pearce, 2013) were met, and missing values were excluded pairwise.

Once two factors were identified, we ran multiple regression to predict the *emotional distress* and *burn-out* from 12 candidate predictors related with gender, age, position, health, care, social support, balance and working conditions. All assumptions of homoscedasticity, normality of residuals, independence of observations, and linearity were found to be met (Pearson’s correlations > 0.7 and Tolerance values > 0.1). There were no outliers in the data and no multicollinearity was present.

## Results

Our results point to a high variability regarding the emotional impact of the pandemic on the researchers in the sample. During lockdown, overall, the participants experienced an intensification of the following emotions, ranked from highest to lowest impact: fatigue (mean = 2.951, SD = 1.287), anxiety (mean = 2.923, SD = 1.168), stress (mean = 2.901, SD = 1.350), frustration (mean = 2.729, SD = 1.283), apathy (mean = 2.599, SD = 1.255), sadness (mean = 2.523, SD = 1.2 = 1.2).

Over 30% of the researcher’s state that stress, fatigue, anxiety, and frustration had increased *quite a lot* or *a lot*. In addition, more than 20% reported significantly increased feelings of sadness, apathy, and helplessness. At the opposite extreme, between 13% and 29% reported no intensification of these emotions, while the majority of researchers reported a small or moderate increase (see Table 2).

**Table 2.** Descriptive statistics and frequencies (n = 1301)

|  | **Anxiety** | **Sadness** | **Apathy** | **Fatigue** | **Frustration** | **Helplessness** | **Stress** |
| --- | --- | --- | --- | --- | --- | --- | --- |
| Average | 2,92 | 2,52 | 2,60 | 2,95 | 2,73 | 2,41 | 2,90 |
| Stand Dev. | 1,17 | 1,13 | 1,26 | 1,29 | 1,28 | 1,23 | 1,35 |
| Nothing | 12,99 | 20,83 | 23,98 | 16,45 | 21,83 | 28,98 | 20,14 |
| Little | 24,14 | 31,59 | 26,21 | 21,68 | 23,83 | 29,21 | 21,29 |
| Moderately | 29,67 | 27,13 | 24,37 | 26,52 | 24,21 | 20,83 | 21,75 |
| Quite a lot | 23,98 | 15,37 | 16,83 | 21,06 | 19,91 | 14,07 | 21,98 |
| A lot | 9,22 | 5,07 | 8,61 | 14,30 | 10,22 | 6,92 | 14,84 |

Exploratory factor analysis showed that the emotional impact during confinement of Covid-19 pandemic on the researchers converged on two main effects. The first factor, which we labelled "emotional distress", consists of a 5-item scale (sadness, helplessness, anxiety, frustration, apathy) and explains 38.9% of the variance with high factorial ranks ranging from 0.60 to 0.84. The second, which corresponds to the syndrome of being overwhelmed, and which we have labelled *burn-out*, is composed of 2 items (tension, fatigue) and explains 20.2% of the variance with factorial ranges also high, ranging from 0.68 to 0.86 (see Table 3).

**Table 3.**
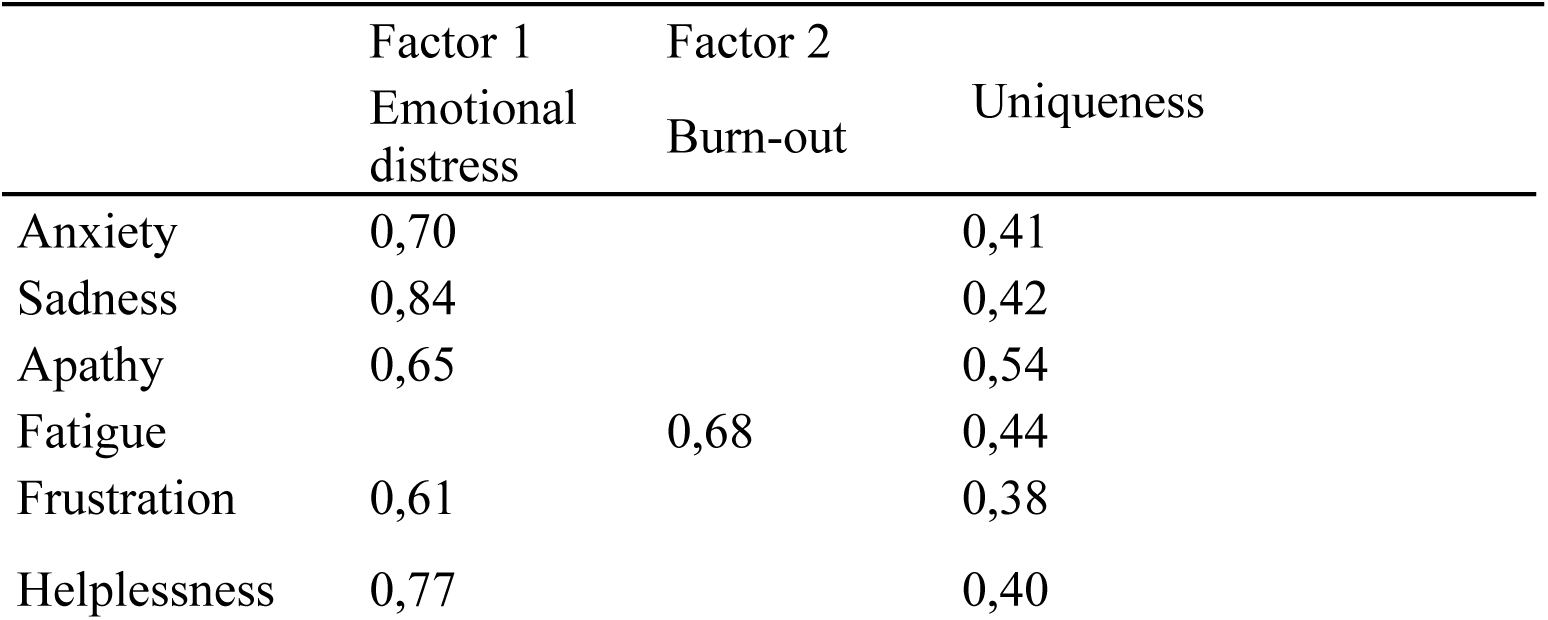
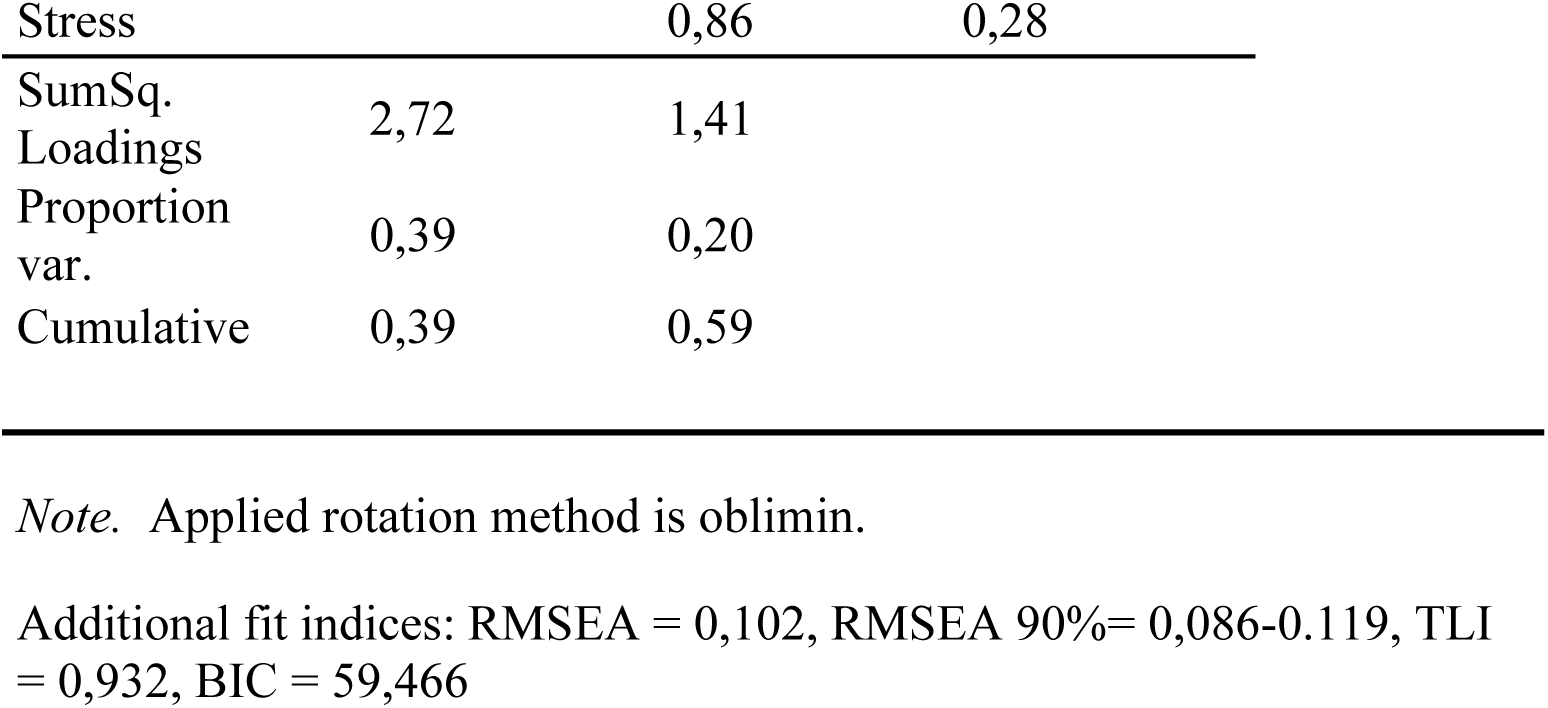
Factor Loadings and characteristics.

Multiple regression analysis showed two models significantly predicting both factors: burn-out (F (12, 1263) = 45.15, p = <, 001) with 10 predictors explaining 29.4% of their variability; and emotional distress (F (12, 126) = 47.18, p = < .001), with eight predictors explaining 30.3% of their variability (see Tables 4 and 5).

**Table 4.** Model Summary of Burn-out and Emotional distress.

|  |  |  |  |  | Durbin-Watson |  |  |
| --- | --- | --- | --- | --- | --- | --- | --- |
| Model | R | R<sup>2</sup> | Adjusted R<sup>2</sup> | RMSE | Autocorrelation | Statistic | p |
| Burn-out | 0.00 | 0.00 | 0.00 | 2.38 | 0.06 | 1.87 | .024 |
|  | 0.55 | 0.30 | 0.29 | 1.99 | -0.02 | 2.04 | .551 |
| Emotional distress | 0.000 | 0.00 | 0.00 | 4.89 | 0.15 | 1.71 | < .001 |
|  | 0.56 | 0.31 | 0.30 | 4.08 | 0.04 | 1.92 | .142 |

**Table 5.** ANOVA of regression models for Burn-out and Emotional distress.

| Model |  | Sum of Squares | df | Mean Square | F | p |
| --- | --- | --- | --- | --- | --- | --- |
| Burn-out ($H_1$) | Regression | 2164.90 | 12 | 180.41 | 45.15 | < .001 |
|  | Residual | 5046.82 | 1263 | 3.99 |  |  |
|  | Total | 7211.72 | 1275 |  |  |  |
| Emotional distress ($H_1$) | Regression | 9423.46 | 12 | 785.29 | 47.18 | < .001 |
|  | Residual | 21022.37 | 1263 | 16.65 |  |  |
|  | Total | 30445.84 | 1275 |  |  |  |

*Note.* The intercept model is omitted, as no meaningful information can be shown.

In relation to burn-out, the variable with the greatest predictive power was the perception that the balance between personal life and research work had been altered and/or damaged during lockdown of Covid-19 pandemic. The other predictors, with decreasing weight, were: *being younger; experiencing health problems; perceiving an worsening current working conditions; caring for others; lack of guidance and direction; feeling professionally isolated; being female; and feeling disconnected from colleagues*. The variables impact of Covid-19 on *future job opportunities* and *research position* were not significantly related (see Table 6).

**Table 6.** Coefficients of burn-out model.

| Explicative variables | Standard values | p |
| --- | --- | --- |
| Isolation or feeling of loneliness | 0.099 | < .001 |
| Disconnection of colleagues | -0.062 | .023 |
| Lack of guidance or supervision | 0.100 | < .001 |
| Current working conditions | -0.104 | < .001 |
| Future job opportunities | 0.023 | .399 |
| Balance with other professional tasks | -0.189 | < .001 |
| Balance with personal life | -0.212 | < .001 |
| Gender | -0.090 | < .001 |
| Age | -0.182 | < .001 |
| Care responsibilities | 0.101 | < .001 |
| Health problems | 0.162 | < .001 |
| Research position | 3.356e -4 | 0.991 |

Regarding emotional distress, the variable with the greatest predictive power was age, whereby younger researchers were more likely to experience emotional distress during lockdown. The other variables that showed a significant relationship with emotional distress were in decreasing weight: *feeling isolated*; *experimenting health problems*; *perceiving a lack of guidance or supervision; holding a junior position; perceiving a worsening in current working conditions* or *future job opportunities*; *difficulties of professional balance*; *being female*; and *difficulties of personal balance*. The variables *feeling disconnected,* and *caregiving* were not significantly related to emotional distress (see Table 7).

**Table 7.**
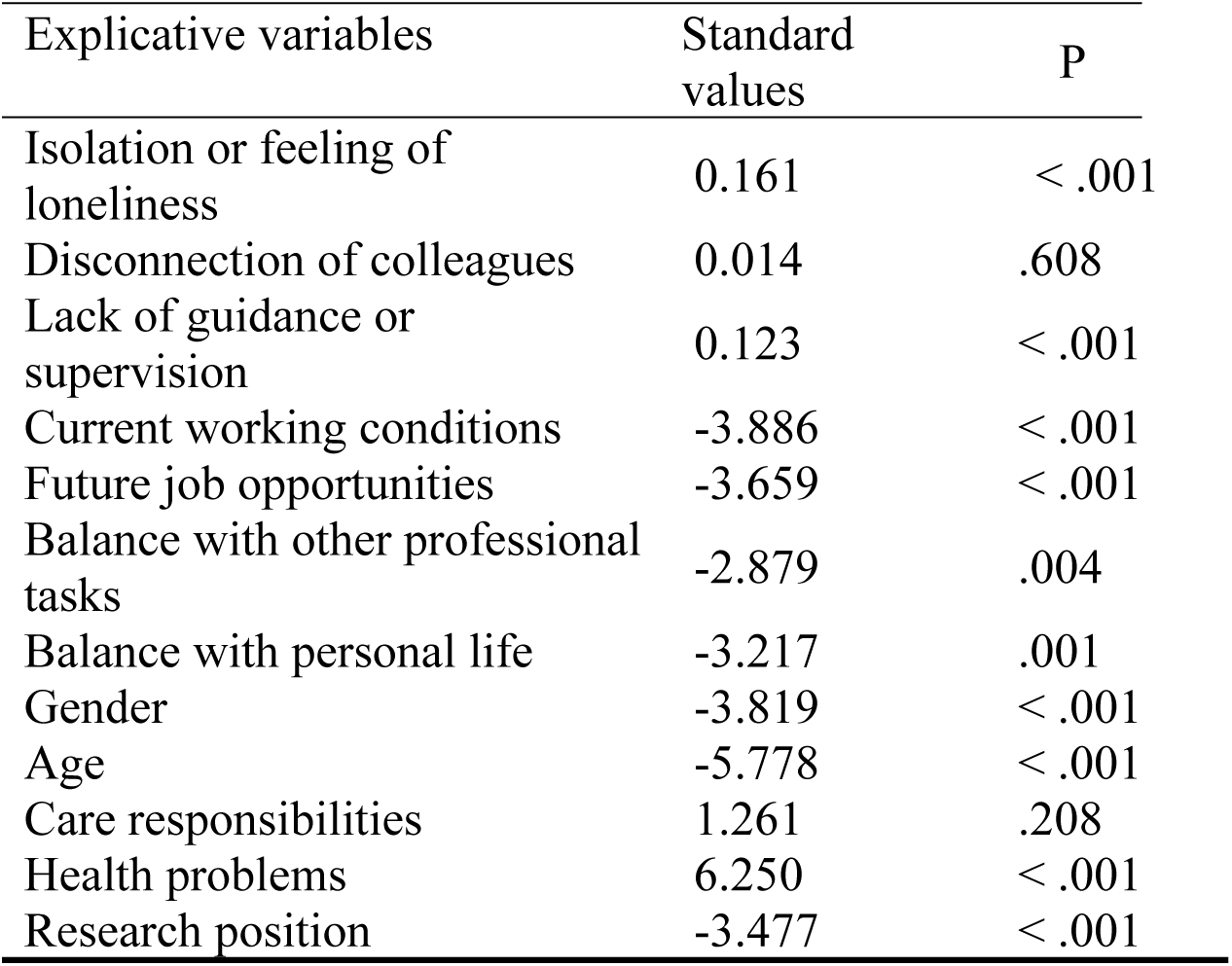
Coefficient of the model of emotional distress.

| Explicative variables | Standard values | P |
| --- | --- | --- |
| Isolation or feeling of loneliness | 0.161 | < .001 |
| Disconnection of colleagues | 0.014 | .608 |
| Lack of guidance or supervision | 0.123 | < .001 |
| Current working conditions | -3.886 | < .001 |
| Future job opportunities | -3.659 | < .001 |
| Balance with other professional tasks | -2.879 | .004 |
| Balance with personal life | -3.217 | .001 |
| Gender | -3.819 | < .001 |
| Age | -5.778 | < .001 |
| Care responsibilities | 1.261 | .208 |
| Health problems | 6.250 | < .001 |
| Research position | -3.477 | < .001 |

## Discussion and conclusions

Our study aimed 1) to analyse the emotional impact of the first confinement during the Covid-19 pandemic on the activity of researchers and 2) to determine the modulating effects on emotional distress of variables such as age, gender, health, research position, working conditions, care responsibilities, health, balance, and social support.

Regarding the first objective, our data reveal a large variability of the effects of the first confinement during the Covid-19 pandemic on the participants’ emotional state. However, it was evident that several negative emotions were intensified for a large majority.

While the intensity experienced also showed high variability, 30% of the participants felt an intense increase in stress, fatigue, and anxiety, and 20% in sadness, apathy, and helplessness. The increase of these negative emotions had as principal effects burnout and emotional distress, impacting the activity of researchers. Psychological stress and exhaustion during the pandemic caused a lack of concentration, focus, and effective development of research activities, as well as increased anxiety levels (Byrom, 2020; Suart et al., 2021).

In the same vein, our results confirm that the conditions of the first confinement exacerbated the usual levels of emotional distress and exhaustion in research activities. Those were already elevated prior to the pandemic (Leonard et al., 2006; Skakni & McAlpine, 2017), increasing the risk of mental health problems for a significant group of researchers.

Beyond these results, the correlation analysis (objective 2) allows us to understand how certain personal conditions and disruptions experienced during the first confinement are predictors of burnout and emotional distress. In this regard, one of the most important predictors in both cases is being young and experiencing health problems. Also, to a lesser extent, a predictor in both cases is to be a woman.

Likewise, the data show that difficulties in reconciling work and family life increased significantly during confinement, leading to an increased risk of burnout, an aspect already noted as usually challenging for researchers in normal conditions (Badri, 2019).

Our data also reveal that the perception of worsening working conditions, lack of social support, and feelings of isolation contributed to increased burnout and emotional distress.

Social support appears to be a protective factor, as lack of guidance and feeling isolated and disconnected is perceived as a significant contributor to both burnout and emotional distress. These data confirm that the feeling of isolation caused by the first confinement is positively associated with higher levels of stress and burnout (Aczel et al., 2021; Gilmartin et al., 2021; Mäkiniemi et al., 2021; Ran, 2018; Sandhu et al., 2021; Sarah et al., 2021; Wang & DeLaquil, 2020).

Such findings align well with previous studies wherein job stress is related to future expectations, overload, role conflict, and peer interaction as risk factors (Hendel & Horn, 2008), which can generate deleterious effects on researchers’ emotional health and productivity (Duxbury et al., 1994).

Furthermore, our data shed light on how the impact of confinement varies for specific populations, pointing to young, early-career female researchers as the most affected.

While studies have already shown that women have higher levels of stress and anxiety (Hankin, et al., 2007; Savitsky, et al., 2020; Suart, Neuman & Truant, 2022) and react differently to them than men (Bekker & van Mens-Verhulst, 2007), our study points out that the emotional effects of first confinement of covid-29 were different in men and women and this was aggravated when they were in the subgroup of younger career researchers, consistently with some recent studies (Suart, Neuman & Truant, 2022). Uncertainty related to the sudden and unexpected transition to distance learning and concern about the future, including academic career assessment procedures (Sahu, 2020), could have played a significant role, with adverse effects on the psychological health of this subgroup of younger individuals.

Gender disparities in the research community are well documented in non-pandemic times and include male-dominated institutional cultures, lack of female mentors, gendered domestic and caregiving responsibilities, and implicit biases in appraisals (Cardel et al., 2020) (e.g., research assignment, peer review outcome, and the number of citations).

A main contribution of the study is clarifying to what extent the factors related to emotional distress intervene differently depending on whether it is burnout or emotional fatigue, which offers complementary explanation on the role of gender and family obligations on the emotional state (Cardel et al., 2020). Surprisingly, caregiving does not directly correlate with emotional distress, although it does with burnout. Although, globally considered, women’s emotional state is worse in our sample, emotional distress is less related to parental or family responsibilities, which suggest further studies are needed to delve on the role of other variables associated with gender in emotional well-being.

Moreover, our data reinforce the the idea that support from peers, colleagues, and leaders is essential to nurture researchers’ well-being and preventing mental health problems (Evans et al., 2019; Loissel, 2020), and offer clear evidence to what extent isolation and lack of emotional and social support might impact on emotional well-being.

In summary, the results of our study affirm that the initial confinement period brought to light certain privileges and exacerbated the emotional repercussions associated with pre-existing inequalities within the research community (Malisch et al., 2020). Furthermore, the COVID-19 pandemic posed a substantial challenge to some of the hard-won gains achieved in gender equality over the past few decades (Matheson, 2020).

It It is worth noting that, prior to the pandemic, concerns about the mental well-being of researchers were already evident. This community operates within a culture that places a premium on productivity at the potential expense of well-being, and it seems to have normalized a pattern of chronic stress and an imbalanced work-life equilibrium (Bartlett et al., 2021; Bekkouche et al., 2021; Guthrie et al., 2018). Their work hours consistently exceed the norm, encompassing increasingly diversified roles and responsibilities within a hypercompetitive job security landscape. Consequently, individuals in managerial roles among researchers may encounter even more pronounced emotional tolls (Kent et al., 2020), although it’s important to note that studies comparing emotional impacts across these various groups are still scarce.

Overall, our findings emphasize the importance of evaluating the concerns and risk perceptions within distinct demographic groups, as these subjective factors can wield considerable influence over their psychological well-being and behavior, including their adherence to appropriate preventive measures (Khosravi et al., 2022; Saita et al., 2021).

Beyond the potential effects of emotional distress in one-off situations such as the first confinement during the Covid-19 pandemic, there is evidence that these effects can be prolonged for months or years, leading to maladaptive behaviours such as substance abuse behaviours, increased chronic depression, and suicide observed three years after the Covid-19 outbreak (Brooks et al., 2020). In the worst cases, this emotional state can lead to psychopathologies such as depression and trauma-related mental health disorders. A recent study by the World Health Organization (2022) confirms the emotional impact of Covid-19 on women and young people, states that anxiety disorders and depression have increased by 25% worldwide and warns of the risk of an emotional pandemic.

Findings underscore the need of offering targeted support to these specific population segments (Matheson, 2020). Additionally, it becomes evident that integrating emotional coping strategies into the training of doctoral students, is imperative (Aitchison & Mowbray, 2013; Sirat, 2012), while also prompting a reconsideration of academic culture itself.

Academic communities should actively cultivate a culture that prioritizes well-being and recognizes the emotional cost of pursuing an academic career (Corbera et al., 2020). This shift in perspective can lead to a more respectful and sustainable academic environment, where personal well-being can be a priority.

Finally, it is paramount to acknowledge the diverse needs, experiences, and vulnerabilities that have been impacted during the Covid-19 pandemic crisis to prevent the chronicization and exacerbation of mental health issues in young academics and learn from them the resilience developed during this period.

Most of our respondents were in Catalonia, so the emotional impact that Covid-19 is having on researchers is restricted to this context. Future work will be needed to expand our understanding of how the pandemic is affecting researchers in different countries, in different types of institutions, at different times in their lives, and with different research conditions, such as available resources, dedication to research, linkage to established research groups, among others. Due to these limitations, the results of the study should not be directly extrapolated to other contexts and be interpreted with caution.

While our findings provide valid information for understanding the emotional impact of Covid-19 on certain populations, it is necessary to delve deeper into the reasons why this impact falls on them. The development of qualitative approaches would be useful to better understand the effects they experienced during the pandemic. Research addressing emotional coping strategies, agency, resilience, and the ability to overcome difficulties could also be useful to understand the diversity of responses in relation to the perceived impact, which would help to identify protective factors, and how differently they relate to gender, age having kids and academic position.

Also, in addition to the long-term negative effects, it would be very useful to further investigate the positive effects and opportunities for researchers arising from the pandemic in order to improve young academic researchers’ development and well-being in a changing and challenging context.

## Footnotes

i The notion of Catalan participants includes those working in the three Catalan-speaking Spanish Autonomous Communities: Valencian Balearic and Catalan

ii The questionnaire is available in Catalan and Spanish and can be found at https://www.researcher-identity.com/researchers-and-covid19

